# Estimating the effect of competition on trait evolution using maximum likelihood inference

**DOI:** 10.1101/023473

**Authors:** Jonathan Drury, Julien Clavel, Marc Manceau, Hélène Morlon

## Abstract

Many classical ecological and evolutionary theoretical frameworks posit that competition between species is an important selective force. For example, in adaptive radiations, resource competition between evolving lineages plays a role in driving phenotypic diversification and exploration of novel ecological space. Nevertheless, current models of trait evolution fit to phylogenies and comparative datasets are not designed to incorporate the effect of competition. The most advanced models in this direction are diversity-dependent models where evolutionary rates depend on lineage diversity. However, these models still treat changes in traits in one branch as independent of the value of traits on other branches, thus ignoring the effect of species similarity on trait evolution. Here, we consider a model where the evolutionary dynamics of traits involved in interspecific interactions are influenced by species similarity in trait values and where we can specify which lineages are in sympatry. We develop a maximum-likelihood based approach to fit this model to combined phylogenetic and phenotypic data. Using simulations, we demonstrate that the approach accurately estimates the simulated parameter values across a broad range of parameter space. Additionally, we develop tools for specifying the biogeographic context in which trait evolution occurs. In order to compare models, we also apply these biogeographic methods to specify which lineages interact sympatrically for two diversity-dependent models. Finally, we fit these various models to morphological data from a classical adaptive radiation (Greater Antillean *Anolis* lizards). We show that models that account for competition and geography perform better than other models. The matching competition model is an important new tool for studying the influence of interspecific interactions, in particular competition, on phenotypic evolution. More generally, it constitutes a step toward a better integration of interspecific interactions in many ecological and evolutionary processes.

---

Interactions between species can be strong selective forces. Indeed, many classical evolutionary theories assume that interspecific competition has large impacts on fitness. Character displacement theory (Brown and Wilson 1956; Grant 1972; Pfennig and Pfennig 2009), for example, posits that interactions between species, whether in ecological or social contexts, drive adaptive changes in phenotypes. Similarly, adaptive radiation theory (Schluter 2000) has been a popular focus of investigators interested in explaining the rapid evolution of phenotypic disparity (Grant and Grant 2002; Losos 2009; Mahler et al. 2013; Weir and Mursleen 2013), and competitive interactions between species in a diversifying clade are a fundamental component of adaptive radiations (Schluter 2000; Losos and Ricklefs 2009; Grant and Grant 2011).

Additionally, social interactions between species, whether in reproductive (Gröning and Hochkirch 2008; Pfennig and Pfennig 2009) or agonistic (Grether et al. 2009, 2013) contexts, are important drivers of changes in signal traits used in social interactions. Several evolutionary hypotheses predict that geographical overlap with closely related taxa should drive divergence in traits used to distinguish between conspecifics and heterospecifics (e.g., traits involved in mate recognition; Wallace 1889; Fisher 1930; Dobzhansky 1940; Mayr 1963; Gröning and Hochkirch 2008; Ord and Stamps 2009; Ord et al. 2011). Moreover, biologists interested in speciation have often argued that interspecific competitive interactions are important drivers of divergence between lineages that ultimately leads to reproductive isolation. Reinforcement, or selection against hybridization (Dobzhansky 1937, 1940), for example, is often thought to be an important phase of speciation (Grant 1999; Coyne and Orr 2004; Rundle and Nosil 2005; Pfennig and Pfennig 2009).

In addition to the importance of interspecific competition in driving phenotypic divergence between species, competitive interactions are also central to many theories of community assembly, which posit that species with similar ecologies exclude each other from the community (Elton 1946). In spite of the importance of interspecific competition to these key ecological and evolutionary theories, the role of competition in driving adaptive divergence and species exclusion from ecological communities has been historically difficult to measure (Losos 2009), because both trait divergence and species exclusion resulting from competition between lineages during their evolutionary history has the effect of eliminating competition between those lineages at the present. Community phylogeneticists have aimed to solve part of this conundrum by analyzing the phylogenetic structure of local communities: assuming that phylogenetic similarity between two species is a good proxy for their ecological similarity, competitive interactions are considered to have been more important in shaping communities comprising phylogenetically (and therefore ecologically) distant species (Webb et al. 2002; Cavender-Bares et al. 2009). However, there is an intrinsic contradiction in this reasoning, because using phylogenetic similarity as a proxy for ecological similarity implicitly (or explicitly) assumes that traits evolved under a Brownian model of trait evolution, meaning that species interactions had no effect on trait divergence (Kraft et al. 2007; Cavender-Bares et al. 2009; Mouquet et al. 2012; Pennell and Harmon 2013).

More generally, and despite the preponderance of classical evolutionary processes that assume that interspecific interactions have important fitness consequences, existing phylogenetic models treat trait evolution within a lineage as independent from traits in other lineages. For example, in the commonly used Brownian motion and Ornstein-Uhlenbeck models of trait evolution (Cavalli-Sforza & Edwards 1967, Felsenstein 1988, Hansen and Martins 1996), once an ancestor splits into two daughter lineages, the trait values in those daughter lineages do not depend on the trait values of sister taxa. Some investigators have indirectly incorporated the influence of interspecific interactions by fitting models where evolutionary rates at a given time depend on the diversity of lineages at that time (e.g., the “diversity-dependent” models of Mahler et al. 2010, Weir and Mursleen 2013). While these models capture some parts of the interspecific processes of central importance to evolutionary theory, such as the influence of ecological opportunity, they do not explicitly account for trait-driven interactions between lineages, as trait values in one lineage do not vary directly as a function of trait values in other evolving lineages.

Recently, Nuismer and Harmon (2015) proposed a model where the evolution of a species’ trait depends on other species’ traits. In particular, they consider a model, which they refer to as the model of phenotype matching, where the probability that an encounter between two individuals has fitness consequences declines as the phenotypes of the individuals become more dissimilar. The consequence of the encounter on fitness can be either negative if the interaction is competitive, resulting in character divergence (matching competition, e.g. resource competition), or positive if the interaction is mutualistic, resulting in character convergence (matching mutualism, e.g. Müllerian mimicry). Applying Lande’s formula (Lande 1976) and given a number of simplifying assumptions—importantly that all lineages evolve in sympatry and that variation in competitors’ phenotypes does not strongly influence the outcome of competition — this model yields a simple prediction for the evolution of a population’s mean phenotype.

Here, we develop inference tools for fitting a simple version of the matching competition model (i.e., the phenotype matching model of Nuismer and Harmon incorporating competitive interactions between lineages) to combined phylogenetic and trait data. We begin by showing how to compute likelihoods associated with this model. Next, we use simulations to explore the statistical properties of maximum likelihood estimation of the matching competition model (parameter estimation as well as model identifiability). While the inclusion of interactions between lineages is an important contribution to quantitative models of trait evolution, applying the matching competition model to an entire clade relies on the assumption that all lineages in the clade are sympatric. However, this assumption will be violated in most empirical cases, so we also developed a method for incorporating data on the biogeographical overlap between species for this model and for the linear and exponential diversity-dependent trait models of Weir & Mursleen (2013), wherein the evolutionary rate at a given time in a tree varies as a function of the number of lineages in the reconstructed phylogeny at that time (see also Mahler et al. 2010).

We then fit the model to data from a classical adaptive radiation: Greater Antillean *Anolis* lizards (Harmon et al. 2003; Losos 2009). Many lines of evidence support the hypothesis that resource competition is responsible for generating divergence between species in both habitat use (e.g., Pacala and Roughgarden 1982) and morphology (Schoener 1970; Williams 1972; see review in Losos 1994). Thus, we can make an *a priori* prediction that model comparison will uncover a signature of competition in morphological traits that vary with habitat and resource use. Given the well-resolved molecular phylogeny (Mahler et al. 2010, 2013) and the relatively simple geographical relationships between species (i.e., many species are restricted to single islands, Rabosky and Glor 2010; Mahler and Ingram 2014), the Greater Antillean *Anolis* lizards provide a good test system for exploring the effect of competition on trait evolution using the matching competition model.

## METHODS

### Likelihood Estimation of the Matching Competition Model

We consider the evolution of a quantitative trait under the matching competition model of Nuismer & Harmon (2015) wherein trait divergence between lineages will be favored by selection. We make the assumption that the outcome of competitive interactions is similar between all members of an evolving clade rather than sensitive to pairwise phenotypic similarity (i.e., that *α* in Eq. 1 of Nuismer and Harmon 2015 is small). This assumption is crucial, as it ensures that the evolution of a population’s mean phenotype is given by a linear model (Eq. S38 in Nuismer and Harmon 2015). Importantly, this implies that the expected distribution of trait values on a given phylogeny follows a multivariate normal distribution (Manceau *et al*., in prep), as is the case for classical models of quantitative trait evolution (Hansen and Martin 1996, Harmon et al. 2010, Weir and Mursleen 2013). In our current treatment of the model, we remove stabilizing selection to focus on the effect of competition (see Discussion). Under these two simplifying assumptions, the mean trait value for lineage *i* after an infinitesimally small time step *dt* is given by (Eq. S38 in Nuismer and Harmon 2015 with *ψ* = 0):

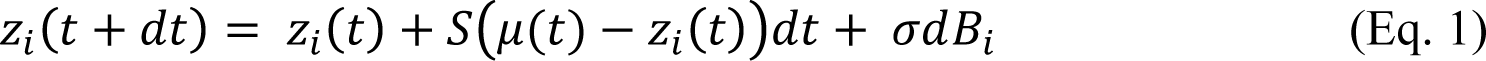

where *z_i_*(*t*) is the mean trait value for lineage *i* at time t, *μ* (*t*) is the mean trait value for the entire clade at time t, *S* measures the strength of interaction (more intense competitive interactions are represented by larger negative values), and drift is incorporated as Brownian motion *σdB_i_* with mean = 0 and variance = *σ*^2^*dt*. Note that when *S* = 0 or *n* = 1 (i.e., when a species is alone), this model reduces to Brownian motion. Under the model specified by Eq. 1, if a species trait value is greater (or smaller) than the trait value average across species in the clade, the species’ trait will evolve towards even larger (or smaller) trait values. We discuss the strengths and limitations of this formulation of the matching competition in the Discussion.

Given that the expected distribution of trait values on a phylogeny under the matching competition model specified in Eq. 1 follows a multivariate normal distribution, it is entirely described with its expected mean vector (made of terms each equal to the character value at the root of the tree) and variance-covariance matrix. Nuismer & Harmon (2015) provide the system of ordinary differential equations describing the evolution of the variance and covariance terms through time (their Eqs.10b and 10c). These differential equations can be integrated numerically from the root to the tips of phylogenies to compute expected variance-covariance matrices for a given set of parameter values and the associated likelihood values given by the multivariate normal distribution.

Additionally, to relax the assumption that all of the lineages in a clade coexist sympatrically, we included a term to specify which lineages co-occur at any given time-point in the phylogeny, which can be inferred, e.g., by biogeographical reconstruction. We define piecewise constant coexistence matrices **A**, where **A***_ij_* equals 1 at time *t* if *i* and *j* are sympatric at that time, and 0 otherwise (Fig. 1). The evolution of the trait value for lineage *i* is then given by:

**Figure 1.**
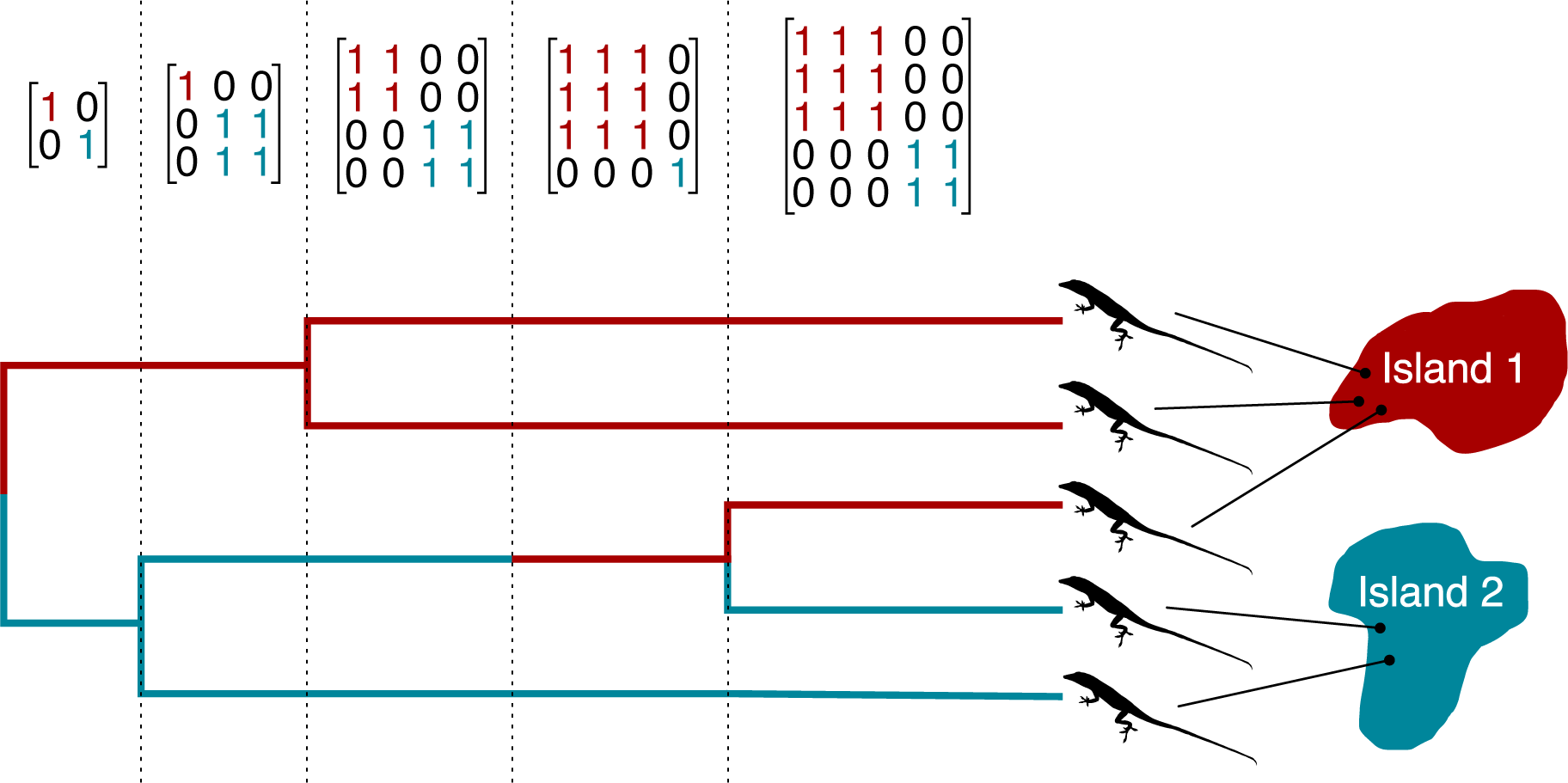
Illustration of geography matrices (defined for each lineage at every node and after each dispersal event inferred, e.g., by stochastic mapping) delineating which lineages interact in sympatry in an imagined phylogeny. These matrices were used to identify potentially interacting lineages for the matching competition and both diversity-dependent models of character evolution (see Eqs. 3-5 in the main text). *Anolis* outline from http://phylopic.org courtesy of Sarah Werning, licensed under Creative Commons. (http://creativecommons.org/licenses/by/3.0/).

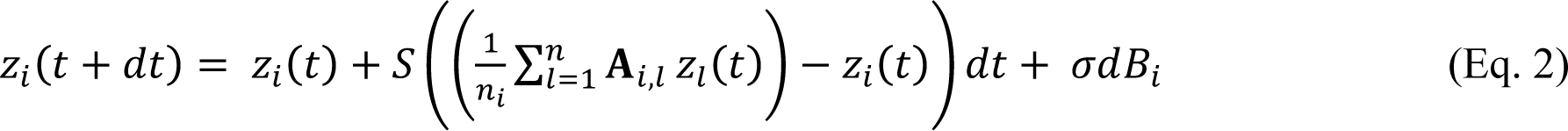

where 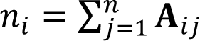 is the number of lineages interacting with lineage *i* at time *t* (equal to the number *n* of lineages in the reconstructed phylogeny at time *t* if all species are sympatric) such that trait evolution is only influenced by sympatric taxa. When a species is alone, **A***_i,i_* = 1, all other **A***_i,j_* = 0, *n_i,i_* = 1, and thus Eq. 2 reduces to the Brownian model.

We show (Appendix S1) that the corresponding system of ordinary differential equations describing the evolution of the variance and covariance terms through time is:

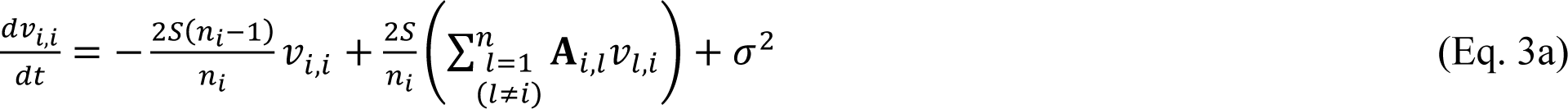

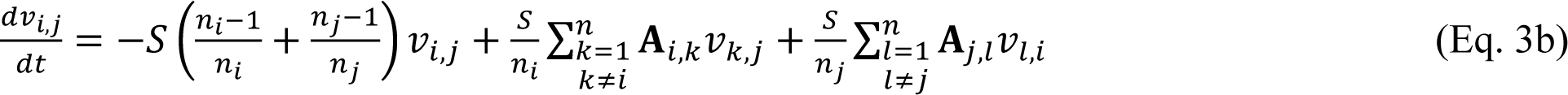

where *v_i,i_* is the variance for each species *i* at time *t* and *v_i,j_* is the covariance for each species pair *i,j* at time *t*. Using numerical integration, we solve this system of ordinary differential equations from the root of the tree to the tips in order to calculate the values of the variance-covariance matrix expected under the model for a given phylogeny and set of parameter values. Specifically, Eq. 3a and 3b dictate the evolution of the variance and covariance values along the branches of the tree; at a given branching event, the variance and covariance values associated to the two daughter species are simply inherited from those of the ancestral species. With the expected variance-covariance matrix at present, we calculate the likelihood for the model using the likelihood function for a multivariate normal distribution (e.g. Harmon et al. 2010). Then, using standard optimization algorithms, we identify the maximum likelihood values for the model parameters. The matching competition model has three free parameters: *σ*^2^, *S* and the ancestral state *z*_0_ at the root. As with other models of trait evolution, the maximum likelihood estimate for the ancestral state is computed through GLS using the estimated variance-covariance matrix (Grafen 1989; Martins and Hansen 1997).

We used the ode function in the R package deSolve (Soetaert et al. 2010) to perform the numerical integration of the differential equations using the “lsoda” solver, and the Nelder-Mead algorithm implemented in the optim function to perform the maximum likelihood optimization. Codes for these analyses are freely available on github (https://github.com/hmorlon/PANDA) and included the R package RPANDA (Morlon et al. 2016).

### Incorporating Geography into Diversity-Dependent Models

Using the same geography matrix **A** described above for the matching competition model (Fig. 1), we modified the diversity-dependent linear and exponential models of Weir & Mursleen (2013) to incorporate biological realism into the models, because ecological opportunity is only relevant within rather than between biogeographical regions. The resulting variance-covariance matrices, **V**, of these models have the elements:

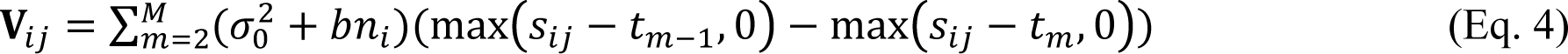

for the diversity-dependent linear model, and

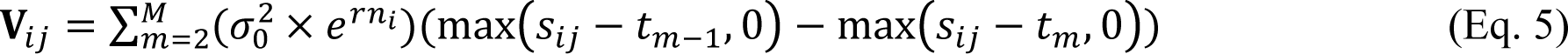

for the diversity-dependent exponential model, where σ_0_^2^ is the rate parameter at the root of the tree, *b* and *r* are the slopes in the linear and exponential models, respectively, *s_ij_* is the shared path length of lineages *i* and *j* from the root of the phylogeny to their common ancestor, *n_i_* is the number of sympatric lineages (as above) between times *t*_*m* -1_ and *t_m_* (where *t*_1_ is 0, the time at the root, and *t*_M_ is the total length of the tree) (Weir & Mursleen 2013). When *b* or *r* = 0, these models reduce to Brownian motion. For the linear version of the model, we constrained the maximum likelihood search such that the term 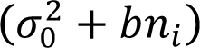 in Eq. 3  0 to prevent the model from having negative evolutionary rates at any *t*_m_.

### Simulation-based Analysis of Statistical Properties of the Matching Competition Model

To verify that the matching competition model can be reliably fit to empirical data, we simulated trait datasets to estimate its statistical properties (i.e., parameter estimation and identifiability using AICc). For all simulations, we began by first generating 100 pure-birth trees using TreeSim (Stadler 2014). To determine the influence of the number of tips in a tree, we ran simulations on trees of size *n* = 20, 50, 100, and 150. We then simulated continuous trait datasets by applying the matching competition model recursively from the root to the tip of each tree (Paradis 2012), following Eq. 1, assuming that all lineages evolved in sympatry. For these simulations, we set *σ*^2^ = 0.05 and systematically varied *S* (-1.5, -1, -0.5, -0.1, or 0). Finally, we fit the matching competition model to these datasets using the ML optimization described above.

To determine the ability of the approach to accurately estimate simulated parameter values, we first compared estimated parameters to the known parameters used to simulate datasets under the matching competition model (*S* and *σ*^2^). We also quantified the robustness of these estimates in the presence of extinction by estimating parameters for datasets simulated on birth-death trees; in addition, we compared the robustness of the matching competition model to extinction to that of the diversity-dependent models. These two latter sets of analyses are described in detail in the Supplementary Appendix 2.

To assess the ability to correctly identify the matching competition model when it is the generating model, we compared the fit (measured by AICc, Burnham and Anderson 2002) of this model to other commonly used trait models on the same data (i.e. data simulated under the matching competition model). Specifically, we compared the matching competition model to (1) Brownian motion (BM), (2) Ornstein-Uhlenbeck/single-stationary peak model (OU, Hansen & Martin 1996), (3) exponential time-dependent (TD_exp_, i.e., the early burst model, or the ACDC model with the rate parameter set to be negative, Blomberg et al. 2003; Harmon et al. 2010), (4) linear time-dependent evolutionary rate (TD_lin_, Weir and Mursleen 2013), (5) linear rate diversity-dependent (DD_lin_, Mahler et al. 2010; Weir and Mursleen 2013), and (6) exponential rate diversity-dependent (DD_exp_, Weir and Mursleen 2013). These models were fitted using geiger (Harmon et al. 2008) when available there (BM, OU, TD_exp_ TD_lin_), or using our own codes, available in RPANDA (Morlon et al. 2016) when they were not available in geiger (DD_lin_, DD_exp_). With the exception of TD_exp_, which we restricted to have decreasing rates through time since the accelerating rates version of the model is unidentifiable from OU (Uyeda et al. 2015), we did not restrict the ML search for the parameters in TD_lin_ or DD models.

We assessed the identifiability of other trait models against the matching competition model by calculating the fit of this model to datasets simulated under the same trait models mentioned above. For BM and OU models, we generated datasets from simulations using parameter values from the appendix of Harmon et al. 2010 scaled to a tree of length 400 (BM, *σ*^2^= 0.03; OU, *σ*^2^ = 0.3, *α* = 0.06). For both the linear and exponential versions of the time- and diversity-dependent models, we simulated datasets with starting rates of *σ*^2^ = 0.6 and ending rates of *σ*^2^ = 0.01, declining with a slope determined by the model and tree (e.g., for time-dependent models, the slope is a function of the total height of the tree; for the TD_exp_ model, these parameters result in a total of 5.9 half-lives elapsing from the root to the tip of the tree, Slater and Pennell 2014). In another set of simulations, we fixed the tree size at 100 tips and varied parameter values to determine the effect of parameter values on identifiability (see Results). As above, we calculated the AICc for all models for each simulated dataset.

Finally, to understand how removing stabilizing selection from the likelihood of the matching competition model affects our inference in the presence of stabilizing selection, we simulated datasets with both matching competition and stabilizing selection on 100 tip trees, across a range of parameter space (*S* = -1, -0.5, and 0, *α* = 0.05, 0.5, and 5, holding *σ*^2^ at 0.05). We fit BM, OU, and matching competition models to these simulated datasets. All simulations were performed using our own codes, available in RPANDA (Morlon 2016).

### Fitting the Matching Competition Model of Trait Evolution to Caribbean *Anolis* Lizards

To determine whether the matching competition model is favored over models that ignore interspecific interactions in an empirical system where competition likely influenced character evolution, we fit the matching competition model to a morphological dataset of adult males from 100 species of Greater Antillean *Anolis* lizards and the time calibrated, maximum clade credibility tree calculated from a Bayesian sample of molecular phylogenies (Mahler et al. 2010, 2013; Mahler and Ingram 2014). We included the first four size-corrected phylogenetic principal components from a set of 11 morphological measurements, collectively accounting for 93% of the cumulative variance explained (see details in Mahler et al. 2013). Each of these axes is readily interprétable as a suite of specific morphological characters (see Discussion), and together, the shape axes quantified by these principal components describe the morphological variation associated with differences between classical ecomorphs in Caribbean anoles (Williams 1972). In addition to the matching competition model, we fit the six previously mentioned models (BM, OU, TD_exp_, TD_lin_, DD_exp_, and DD_lin_) separately to each phylogenetic PC axis in the *Anolis* dataset.

For the matching competition model and diversity-dependent models, to determine the influence of uncertainty in designating clades as sympatric and allopatric, we fit the model for each trait using 101 sets of geography matrices (i.e., **A** in Eq. 1b, 2, & 3, see Fig. 1): one where all lineages were set as sympatric, and the remaining 100 with biogeographical reconstructions from the output of the make.simmap function in phytools (Revell 2012). To simplify the ML optimization, we restricted *S* to take negative values while fitting the matching competition model including the biogeographical relationships among taxa (i.e., we forced the optimization algorithm to only propose *S* values  <u>0</u>).

## RESULTS

### Statistical Properties of the Matching Competition Model

Across a range of *S* values, maximum likelihood optimization returns reliable estimates of parameter values for the matching competition model (Fig. 2). As the number of tips increases, so does the reliability of maximum likelihood parameter values (Fig. 2). Parameter estimates remain reliable in the presence of extinction, unless the extinction fraction is very large (i.e.,  0.6; Supplementary Appendix 2). When datasets are simulated under the matching competition model, model selection using AICc generally picks the matching competition model as the best model (Figs. 3, S1); the strength of this discrimination depends on both the *S* value used to simulate the data and the size of the tree (Figs. 3, S1). For example, when *S* = -0.1, the matching competition model often has a higher AICc than Brownian motion, largely due to the fact that the Brownian motion model has one less parameter.

**Figure 2.**
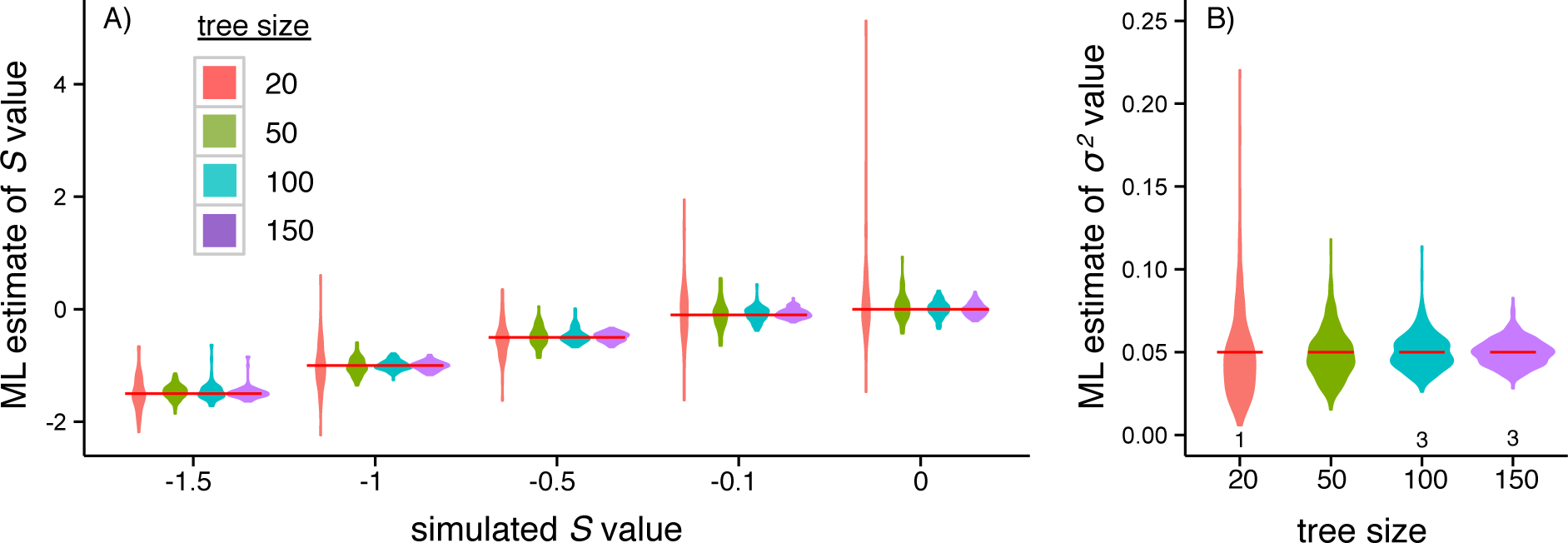
Parameter estimation under the matching competition model. As tree size increases and/or the magnitude of competition increases (i.e., the *S* parameter in the matching competition model becomes more negative), so does the accuracy of ML parameter estimates of (A) *S* (*n* = 100 for each tree size and *S* value combination; red horizontal lines indicate the simulated *S* value) and (Β) σ^2^ (*n* = 500 for each tree size; red horizontal lines indicate the simulated value). In a small number of cases (7/2000), the ML estimate for *σ^2^* was unusually large ( > 0.25), and we removed these rare cases for plotting. The numbers below the violin plots in (B) show the number of outliers removed for each tree size.

**Figure 3.**
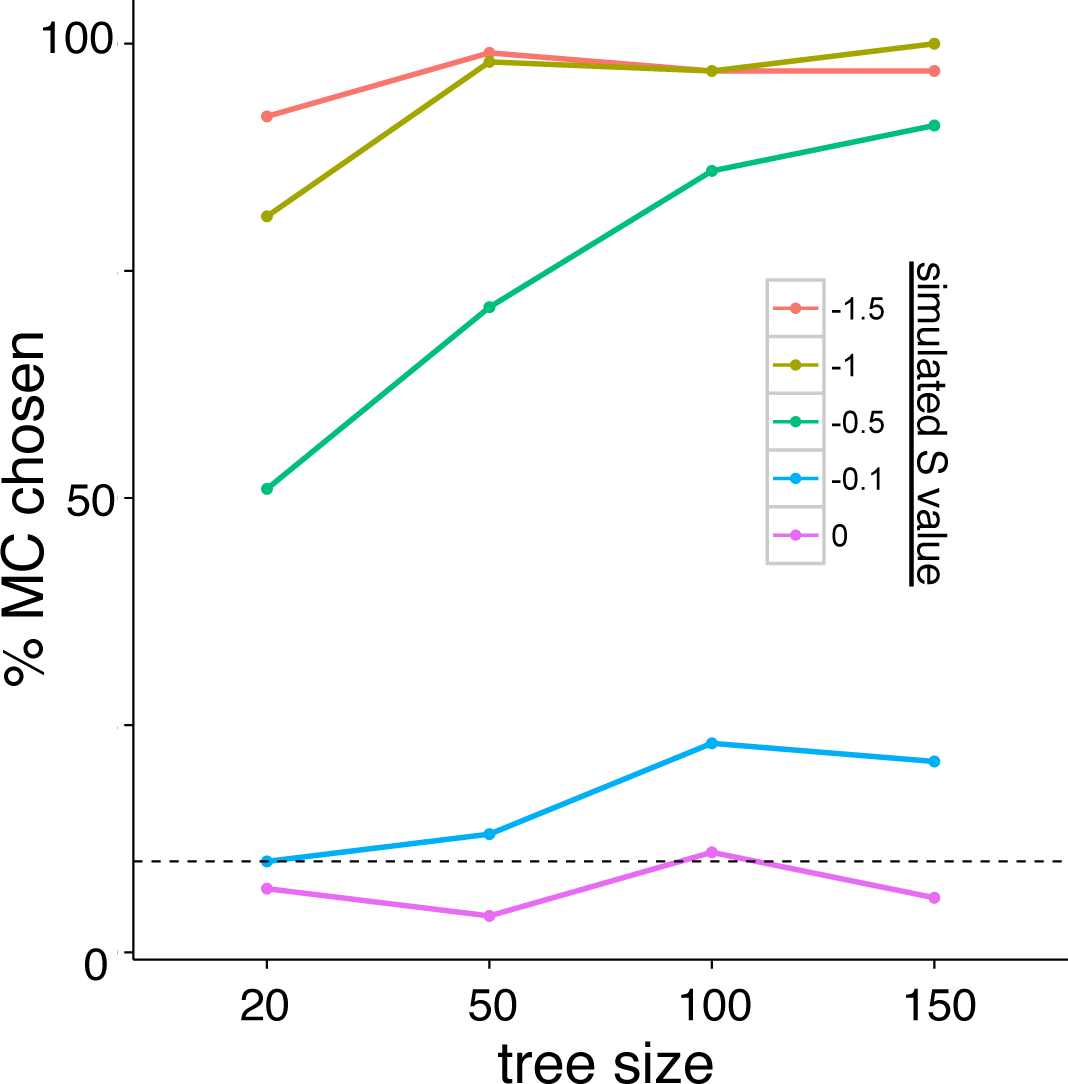
AICc support for datasets simulated under the matching competition (MC) model increases with tree size and with increasing levels of competition (i.e., increasingly negative *S* values). The dotted line denotes 10%.

Simulating datasets under BM, OU, DD_exp_, and DD_lin_ generating models, we found that in most scenarios, and in most parameter space, these models are distinguishable from the matching competition model (Fig. 4a,b,e,f, Fig. S2). As with the matching competition model, the ability to distinguish between models using AICc generally increases with increasing tree sizes (Fig. 4) and with increasing magnitude of parameter values (Fig. S2). When character data were simulated under a TD_lin_ model of evolution, the matching competition and/or the diversity-dependent models tended to have lower AICc values than the TD_lin_ model, especially among smaller trees (Figure 4d). For data generated under a TD_exp_ model, model selection always favored the matching competition model over the TD_exp_ model (Fig. 4c).

Though the current implementation of the maximum likelihood tools for the matching competition do not incorporate stabilizing selection, simulating datasets with both matching competition and stabilizing selection reveals that as the strength of stabilizing selection increases relative to the strength of competition (i.e., α as increases relative to *S*), AICc model selection shifts from favoring the matching competition model (under large S, small α scenarios) to favoring the OU model (under small S, large *α* scenarios) (Fig. S3). Likewise, maximum likelihood increasingly underestimates the value of *S* as the value of *α* increases (Fig. S4).

### Competition in Greater Antillean *Anolis* Lizards

For the first four phylogenetic principal components describing variation in *Anolis* morphology, we found that models that incorporate species interactions fit the data better than models that ignore them (Table 1). PC1, which describes variation in hindlimb/hindtoe length (Mahler et al. 2013), is fit best by the matching competition model. PC2, which describes variation in body size (snout vent length) is fit best by the linear diversity-dependent model. PC3, which describes variation in forelimb/foretoe length, and PC4, which describes variation in lamellae number are fit with mixed support across the models included, but with models incorporating species interactions providing the best overall fits.

Additionally, for every PC axis, the best-fit models were ones that incorporated the geographic relationships among species in the tree, and these conclusions were robust to uncertainty in ancestral reconstructions of sympatry (Table 1).

**Figure 4.**
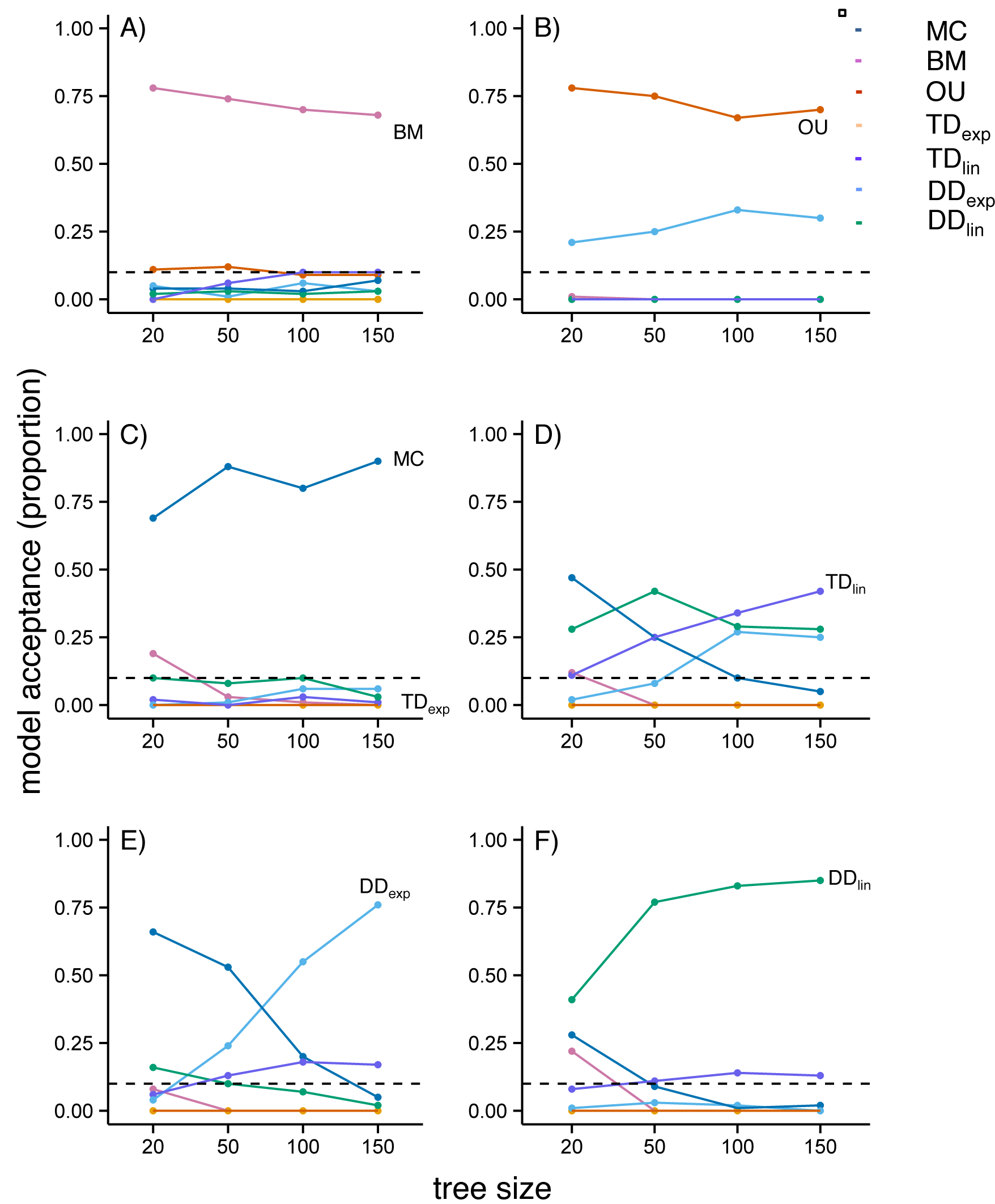
Identifiability simulation results for the matching competition (MC) model. When the generating model is either (A) BM, (B) OU, (E) DD_exp_ (for larger trees) or (F) DD_lin_, the generating model is largely favored by model selection. However, both (C) TD_exp_ and (D) TD_lin_ (for smaller trees) are erroneously rejected as the generating model. The dotted lines denote 10%.

**Table 1.**
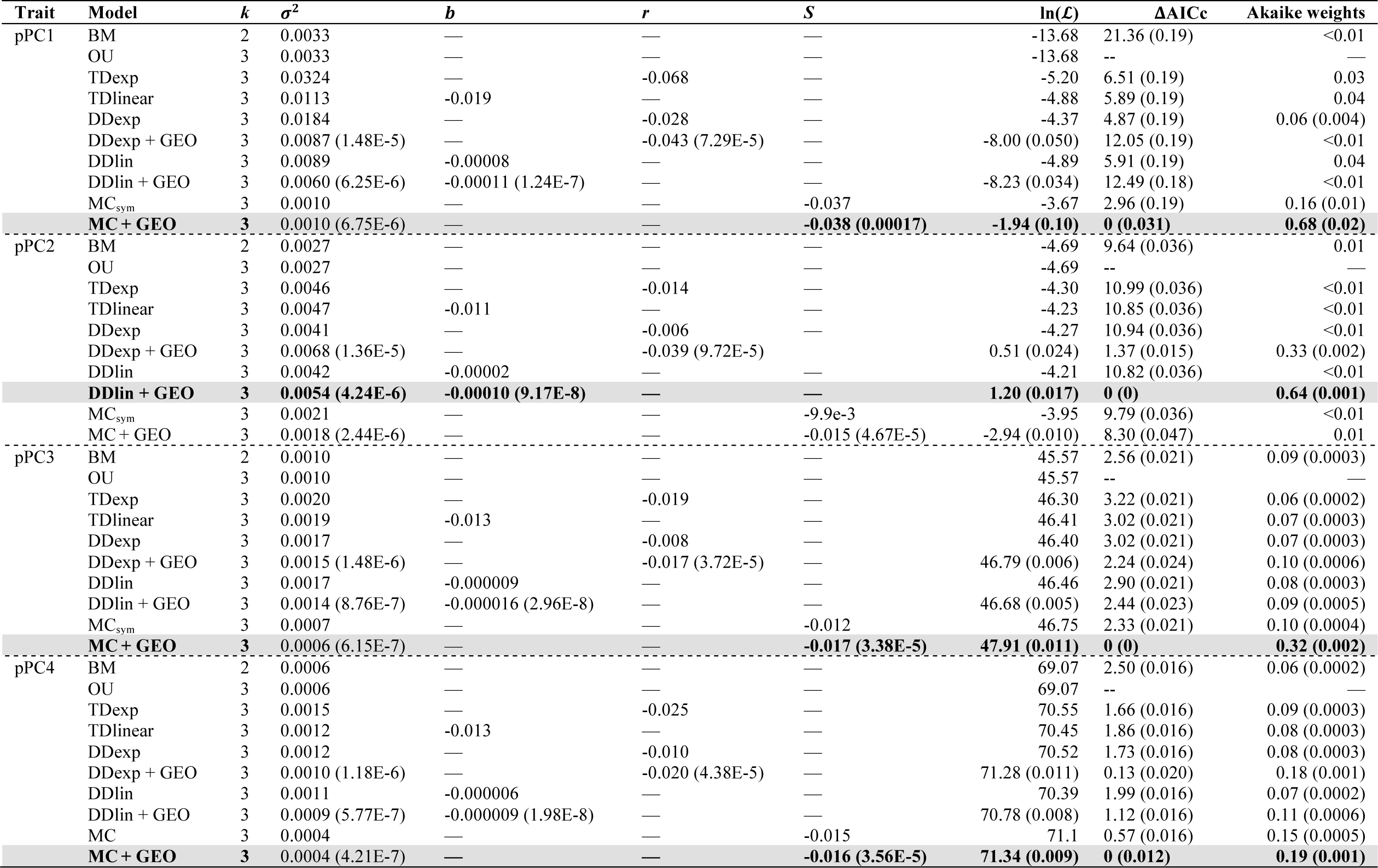
Comparison of model fits for the first four phylogenetic principal components of a morphological dataset of Greater Antillean anoles. Models run incorporating geography matrices are indicated by “+ GEO”, and models with the lowest AICc for each trait are shaded and written in bold text. Parameter values presented follow the nomenclature of Eqs. 2-4 in the main text, and *k* represents the number of parameters estimated for each model. Note that TD_exp_ is the ACDC model (or the early-burst model when *r* Δ 0). OU model weights were excluded because the ML estimates of *α* equaled 0 for all PC axes, and thus the OU model was equivalent to BM. Median (standard error) of parameter estimates, ΔAICc values, and Akaike weights are presented for fits across 100 sampled stochastic maps of *Anolis* biogeography (standard errors are omitted for Akaike weights < 0.05).

## DISCUSSION

The inference methods we present here represent an important new addition to the comparative trait analysis toolkit. Whereas previous models had not accounted for the influence of trait values in other lineages on character evolution, the matching competition model takes these into account. Furthermore, extending both the matching competition model and two diversity-dependent trait evolution models to incorporate geographic networks of sympatry further extends the utility and biological realism of these models.

We found that the matching competition model has increasing AICc support and accuracy of parameter estimation with increasing tree sizes and competition strength. We also found that, for most of the generating models we tested, AICc-based model selection does not tend to erroneously select the matching competition model (i.e., these models are identifiable from the matching competition model). As with all other models, the statistical properties of the matching competition model will depend on the size and shape of a particular phylogeny as well as specific model parameter values. Future investigators can employ other approaches, such as phylogenetic Monte Carlo and posterior predictive simulations directly on their empirical trees (Boettiger et al. 2012, Slater & Pennell 2014), to assess the confidence they can have in their results.

We did, however, find that data generated under time-dependent models were often fit better by models that incorporate interspecific interactions (i.e., density-dependent and matching competition models) (Fig. 4c,d). This was especially true for the TD_exp_ model, often referred to as the early-burst model—the matching competition model nearly always fit data generated under the TD_exp_ model better than the TD_exp_ model (Fig. 4c). We do not view this as a major limitation of the model for two reasons. First, the TD_exp_ model is known to be statistically difficult to estimate on neontological data alone (Harmon et al. 2010; Slater et al. 2012a; Slater and Pennell 2014). Secondly, and more importantly, time-dependent models are not process-based models, but rather incorporate time since the root of a tree as a proxy for ecological opportunity or available niche space (Harmon et al. 2010; Mahler et al. 2010; Slater 2015). The matching competition and density-dependent models explicitly account for the interspecific competitive interactions that time-dependent models purport to model, thus we argue that these process-based models are more biologically meaningful than time-dependent models (Moen and Morlon 2014).

We did not incorporate stabilizing selection in our model. Preliminary analyses suggested that *S* and ( are not identifiable (though their sum may be), as competition and stabilizing selection operate in opposite directions. As a result, when trait data are simulated with simultaneous stabilizing selection and matching competition, the strength of competition is underestimated. In addition, which model is chosen by model selection depends on the ratio of the strength of attraction toward an optimum to the strength of competition, with Brownian model being selected at equal strengths (Figs. S3, S4). Given that many traits involved in competitive interactions are also likely to have been subject to stabilizing selection (i.e., extreme trait values eventually become targeted by negative selection), statistical inference under the matching competition model without stabilizing selection is likely to underestimate the true effect of competition on trait evolution. Future work aimed at directly incorporating stabilizing selection in the inference tool could provide a more accurate quantification of the effect of competition, although dealing with the non-identifiability issue may require incorporating additional data such as fossils.

Because the matching competition model depends on the mean trait values in an evolving clade, maximum likelihood estimation is robust to extinction, whereas the diversity-dependent models are less so (Appendix S2, Figs. S5-S8). Nevertheless, given the failure of maximum likelihood to recover accurate parameter estimates of the matching competition model at high levels of extinction (*μ: λ*  0.6), we suggest that these models should not be used in clades where the extinction rate is known to be particularly high. In such cases, it would be preferable to modify the inference framework presented here to include data from fossil lineages (Slater et al. 2012a) by adapting the ordinary differential equations described in Eq. 3a and 3b for non-ultrametric trees.

For all of the traits we analyzed, we found that models incorporating both the influence of other lineages and the specific geographical relationships among lineages were the most strongly supported models (though less strikingly for PC3 and PC4). Incorporating uncertainty in biogeographical reconstruction, which we encourage future investigators to do in general, demonstrated that these conclusions were robust to variation in the designation of allopatry and sympatry throughout the clade. The matching competition model is favored in the phylogenetic principal component axis describing variation in relative hindlimb size. Previous research demonstrates that limb morphology explains between-ecomorph variation in locomotive capabilities and perch characteristics (Losos 1990, 2009; Irschick et al. 1997), and our results suggest that the evolutionary dynamics of these traits have been influenced by the evolution of limb morphology in other sympatric lineages. These results support the assumption that interspecific interactions resulting from similarity in trait values are important components of adaptive radiations (Losos 1994, Schluter 2000), a prediction that has been historically difficult to test (Losos 2009, but see Mahler et al. 2010). In combination with previous research demonstrating a set of convergent adaptive peaks in morphospace to which lineages are attracted (Mahler et al. 2013), our results suggest that competition likely played an important role in driving lineages toward these distinct peaks. Because we expect the presence of selection toward optima to lead to underestimation of the *S* parameter in the matching competition model (Figs. S3, S4), we would have likely detected an even stronger effect of competition in the *Anolis* dataset if we had included stabilizing selection. Recently, Uyeda and colleagues (2015) demonstrated that the use of principal components can bias inferences of trait evolution. We used BM-based phylogenetic PC axes here, which should reduce this potential bias (Revell 2009). We recognize that there is some circularity in assuming BM in order to compute phylogenetic PC axes before fitting other trait models to these axes; a general solution to address this circularity problem remains to be found (Uyeda et al. 2015). Uyeda & colleagues suggested that using phylogenetic PC axes sorts the traits according to specific models. In the Greater Antillean *Anolis* lizards, the first axes are easily interpretable as specific suites of traits relevant to competitive interactions, and our results suggest that competition played an important role in shaping the evolution of these traits.

The linear version of Nuismer & Harmon’s (2015) model (Eq. 1) results from making the simplifying assumption that the outcome of competition is not highly sensitive to variation in sympatric competitors’ phenotypes (i.e., that *α* in their Eq. 1, and as a result also *S* in our equations, are small). We used this version here, since currently available likelihood tools for trait evolution rely on the multivariate normal distribution, which is to be expected only for this linear form of the model. The current formulation (Eq. 1) corresponds to a scenario in which the rate of phenotypic evolution in a lineage gets higher as the lineage deviates from the mean phenotype, although character displacement theory, for example, posits that selection for divergence should be the strongest when species are most ecologically similar (Brown and Wilson 1956). Given this formulation of the model, large *S* values are not to be expected, and we indeed found relatively small *S* values when fitting the model to the *Anolis* dataset. Investigators finding high *S* values should treat them with caution and consider enforcing bounds on the likelihood search. Nevertheless, the developments presented here provide an important new set of tools for investigating the impact of interspecific interactions on trait evolution, and researchers can perform posterior simulations to assess the realism of the resulting inference. Future development of likelihood-free methods, such as Approximate Bayesian Computation (Slater et al. 2012b; Kutsukake and Innan 2013), may be possible for fitting the version of the model in which the outcome of competitive interactions depends on distance in trait space.

We imagine that the matching competition model and biogeographical implementations of diversity-dependent models will play a substantial role in the study of interspecific competition. For example, by comparing the fits of the matching competition model with other models that do not include competitive interactions between lineages, biologists can directly test hypotheses that make predictions about the role of interspecific interactions in driving trait evolution. In other words, while the effect of competition has been historically difficult to detect (Losos 2009), it may be detectable in the contemporary distribution of trait values and their covariance structure (Hansen and Martins 1996; Nuismer and Harmon 2015). The ability to consider trait distributions among species that arise from a model explicitly accounting for the effect of species interactions on trait divergence is also an important step toward a more coherent integration of macroevolutionary models of phenotypic evolution in community ecology.

There are many possible extensions of the tools developed in this paper. In the future, empirical applications of the model can be implemented with more complex geography matrices that are more realistic for mainland taxa (e.g., using ancestral biogeographical reconstruction, Ronquist and Sanmartín 2011; Landis et al. 2013), and can also specify degrees of sympatric overlap (i.e., syntopy). Additionally, the current version of the model is rather computationally expensive with larger trees (on a Mac laptop with a 2.6 GHz processor, maximum likelihood optimization for the matching competition model takes several minutes for a tree with 50 tips and can take 30 minutes or longer on 100 tip trees). Further work developing an analytical solution to the model may greatly speed up the likelihood calculation and permit the inclusion of stabilizing selection.

The current form of the model assumes that the degree of competition is equal for all interacting lineages. Future modifications of the model, such as applications of stepwise AICc algorithms (Alfaro et al. 2009; Thomas and Freckleton 2012; Mahler et al. 2013) or reversible-jump Markov Chain Monte Carlo (Pagel and Meade 2006; Eastman et al. 2011; Rabosky 2014; Uyeda and Harmon 2014), may be useful to either identify more intensely competing lineages or test specific hypotheses about the strength of competition between specific taxa. Improvements could also be made on the formulation itself of the evolution of a species’ trait as a response to the phenotypic landscape in which the species occurs. Moreover, a great array of extensions will come from modeling species interactions not only within clades, but also among interacting clades, as in the case of coevolution in bipartite mutualistic or antagonistic networks, such as plant-pollinator or plant-herbivore systems.

## Acknowledgements

We thank J. Weir for providing R code for diversity-dependent models and E. Lewitus, O. Missa, F. Anderson, L. Harmon, and two anonymous reviewers for helpful comments on the manuscript. This research was funded by the Agence Nationale de la Recherche (grant CHEX-ECOEVOBIO) and the European Research Council (grant 616419-PANDA) to HM.

## Supplementary Material

### *Supplementary Appendix 2:* Estimating the effect of extinction on parameter estimation for the matching competition and density-dependent models

Given that the matching competition and diversity-dependent models take into account the number of interacting lineages, extinction may affect our ability to recover true parameter values. To estimate the impact of extinction, we simulated 100 trees with 100 extant species, varying the extinction fraction (*μ*:(= 0.2, 0.4, 0.6, and 0.8). As above, we recursively simulated traits using the matching competition model with *σ*^2^ = 0.05 and *S* = -1.5, -1, -0.5, -0.1, or 0, and the linear and exponential diversity-dependent models with starting rates of *σ*^2^ = 0.6 and ending rates of *σ*^2^ = 0.01. We then estimated the maximum likelihood parameter estimates for the generating models by fitting the models to the trait values for extant species and the tree with extinct lineages removed. In the case of the matching competition model, because many simulated birth-death trees with high extinction rates have substantially older root ages, the simulated trait datasets for some trees had very large variances. For these biologically unrealistic trait datasets (i.e., variance in trait values  1×10^8^), ML does not yield reliable parameter estimates, so we removed them from further analyses (the sample size of included simulations is reported in Fig. S5, S6).

Parameter estimates are quite robust to extinction under the matching competition model (Fig. S5, S6), and much more so than under both diversity-dependent models (Fig. S7, S8). Under the matching competition model, the maximum likelihood optimization returns reliable estimates of *S* and *σ*^2^ values used to simulate datasets on trees with extinct lineages (Fig. S5, S6), although the estimates become much less reliable with larger extinction fractions, likely because simulations under the matching competition model were unbounded, resulting in trait datasets with biologically unrealistic variances. Under both diversity-dependent models, the magnitude of both the slope and *σ*^2^ parameter values are increasingly underestimated with increasing extinction fractions (Fig. S7, S8).

**Supplementary Figure 1.**
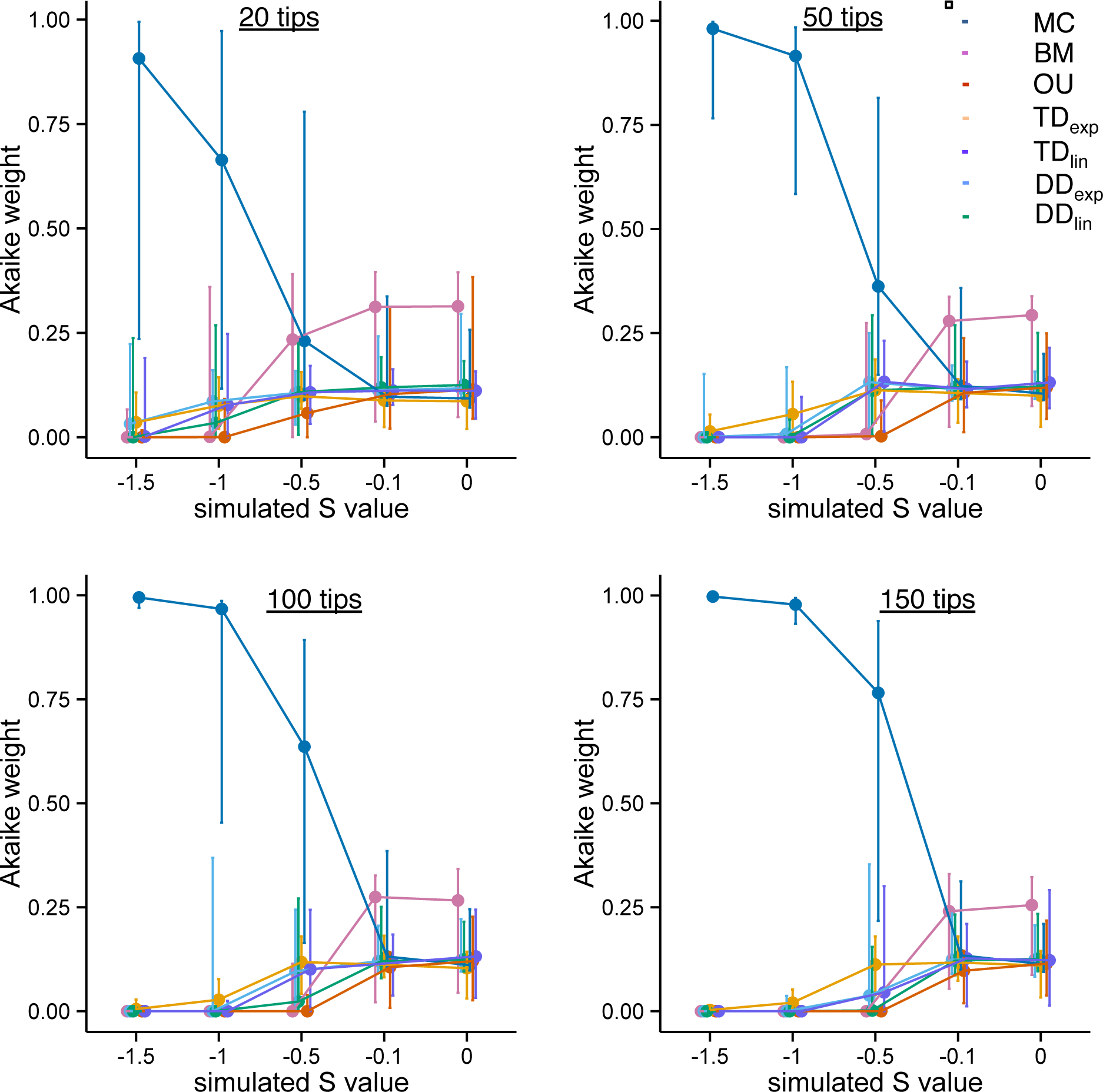
As tree size and/or the degree of competition (*S*) increases, model selection becomes more reliable. Comparison of Akaike weights (median & 90% CIs) for NH, BM, OU, and EB models when simulated under various levels of competition (*S* = -1.5, -1, -0.5, -0.1, and 0) for trees with 20, 50, 100, and 150 tips.

**Supplementary Figure 2.**
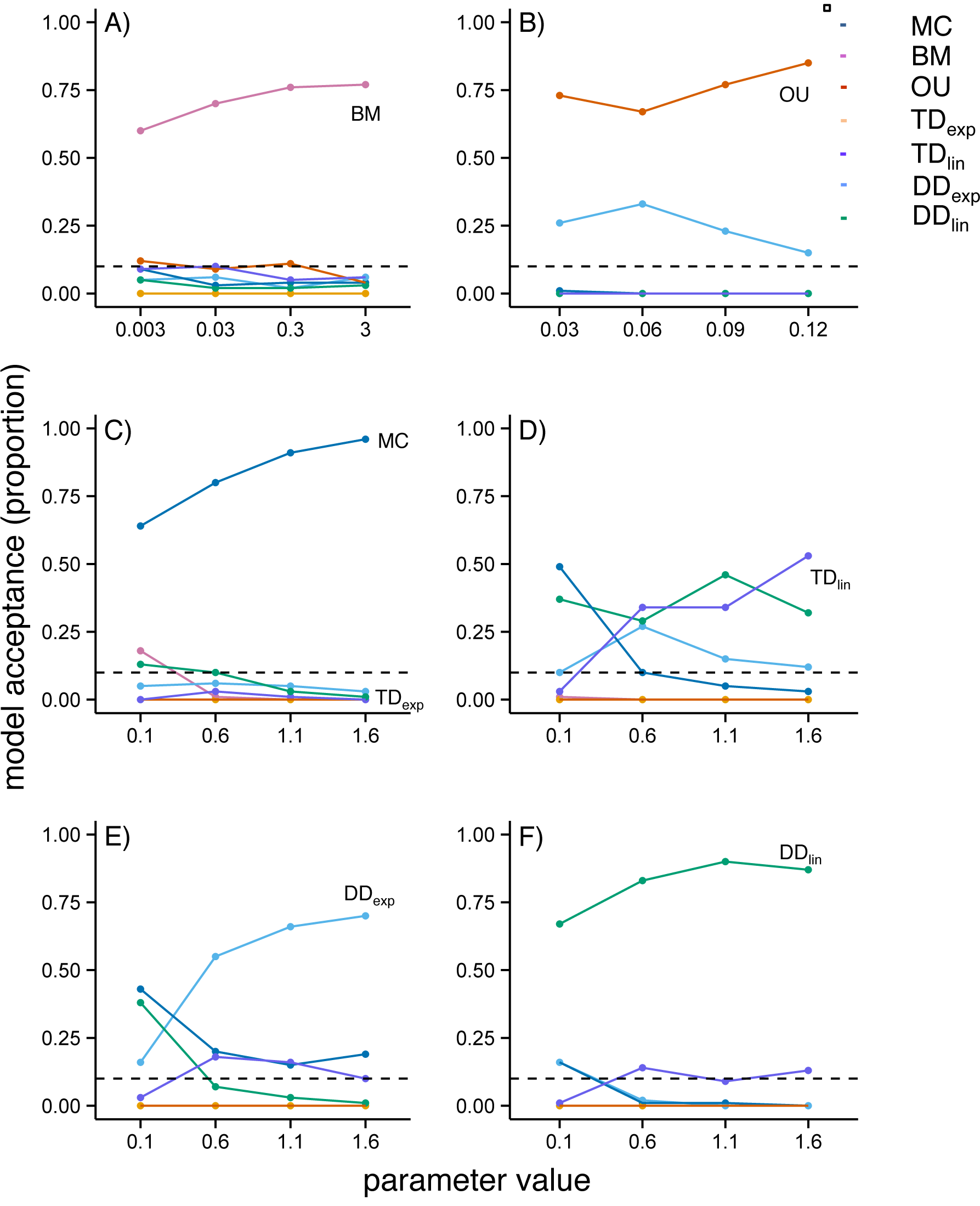
Identifiability simulation results for the matching competition model as a function of varying parameter values of the generating models. Parameter values are (A) *σ*^2^ for BM, (B) *α* for OU (*σ*^2^ was fixed at 0.3), and the *σ*^2^ value at the root for (C) TD_exp_, (D) TD_lin_, (E) DD_exp_, and (F) DD_lin_ (for C-F, *σ*^2^ at the present was fixed at 0.01).

**Supplementary Figure 3.**
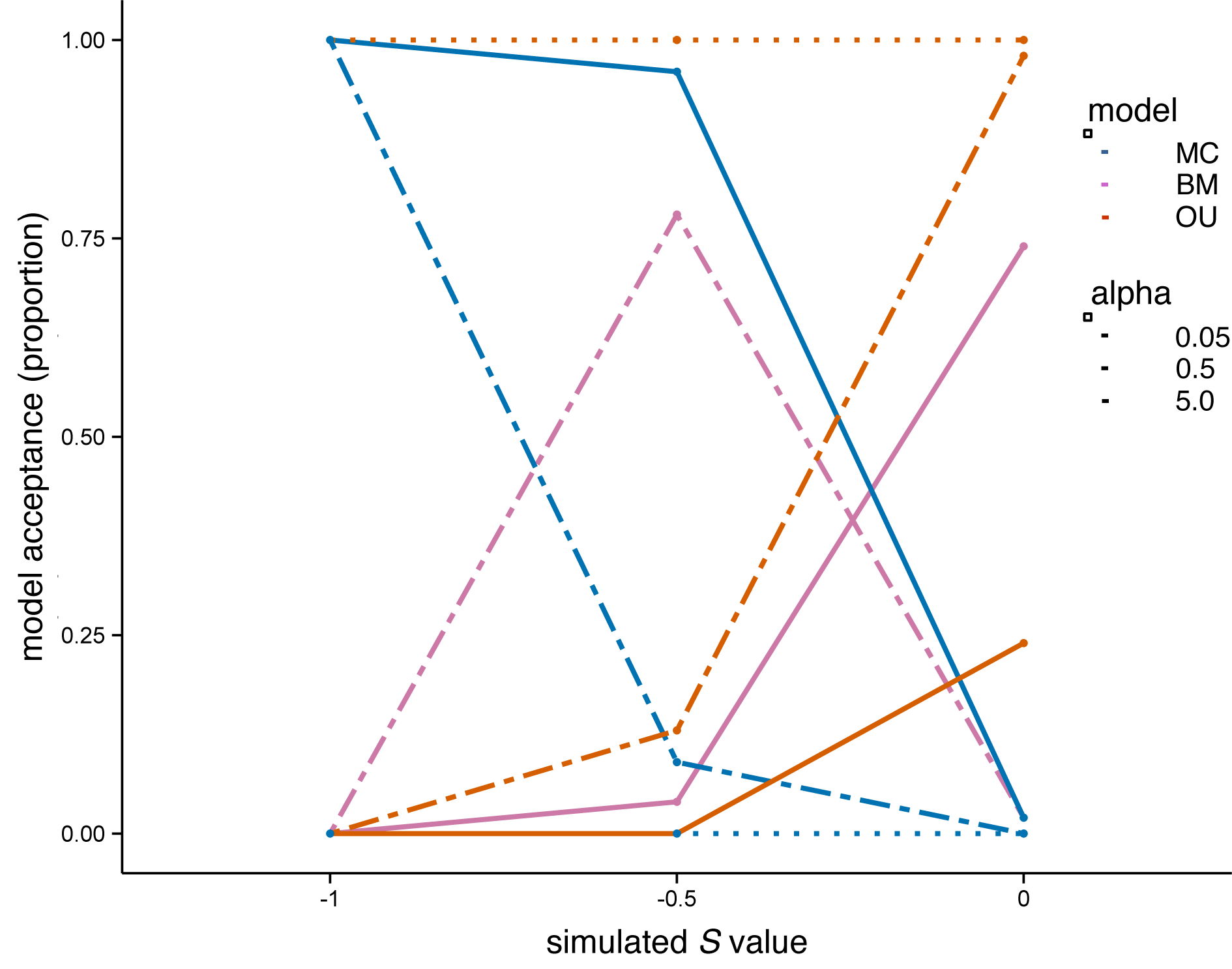
The effect of incorporating stabilizing selection into trait evolution on model selection. For datasets generated under the matching competition model with stabilizing selection included, as the ratio of the strength competition (*S*) to the strength of selection toward an optimum (*α*) varies, so does the model preferred by model selection.

**Supplementary Figure 4.**
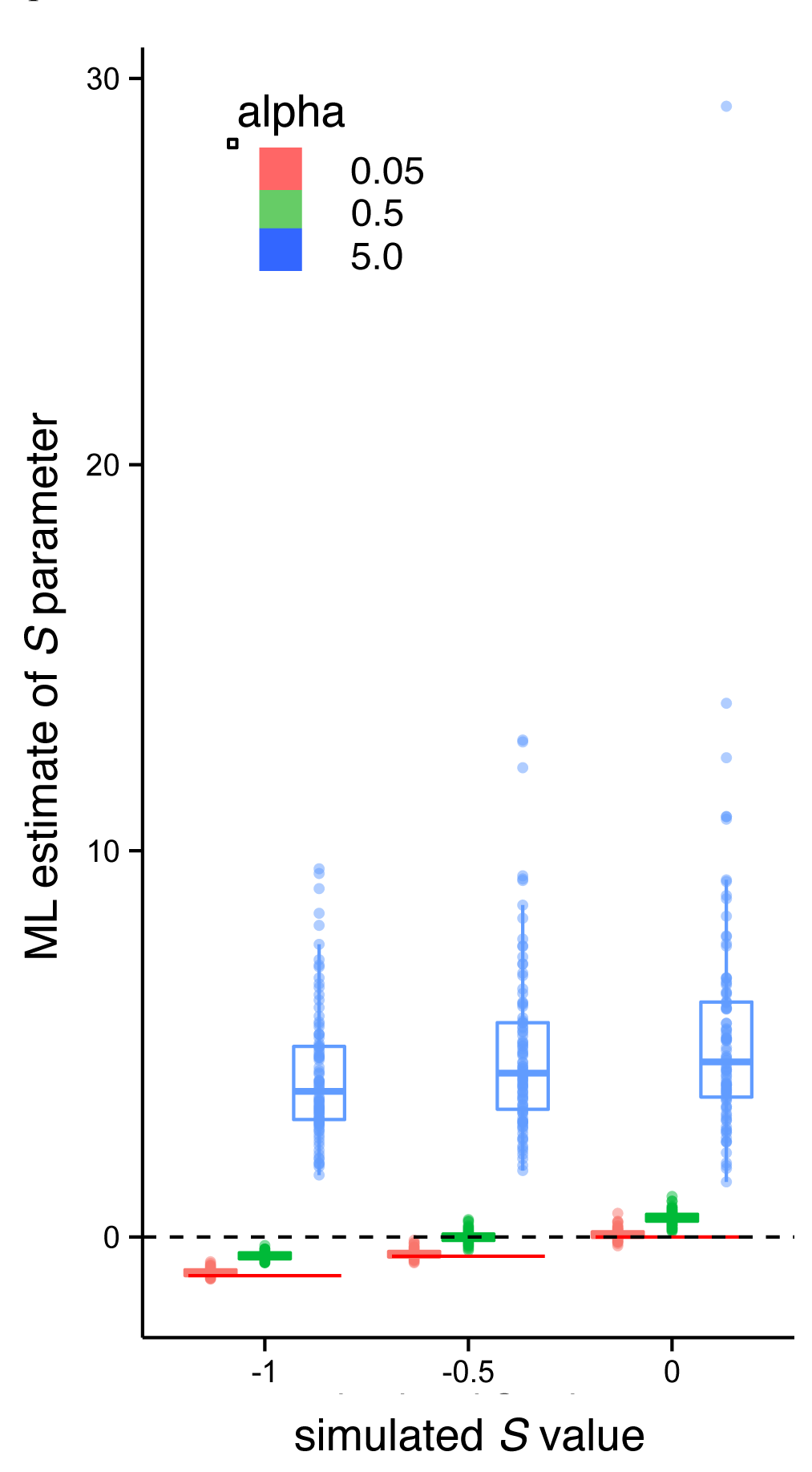
The effect of incorporating stabilizing selection into trait evolution on parameter estimation. As the strength of stabilizing selection increases (i.e., as *α* increases), maximum likelihood under the matching competition model underestimates the true *S* value used to simulate datasets. Positive *S* values represent selection toward, rather than away, from the clade mean and are thus expected when the ratio of *α* to *S* is large. The horizontal red line represents the simulated *S* value, and the dashed horizontal line represents *S* = 0.

**Supplementary Figure 5.**
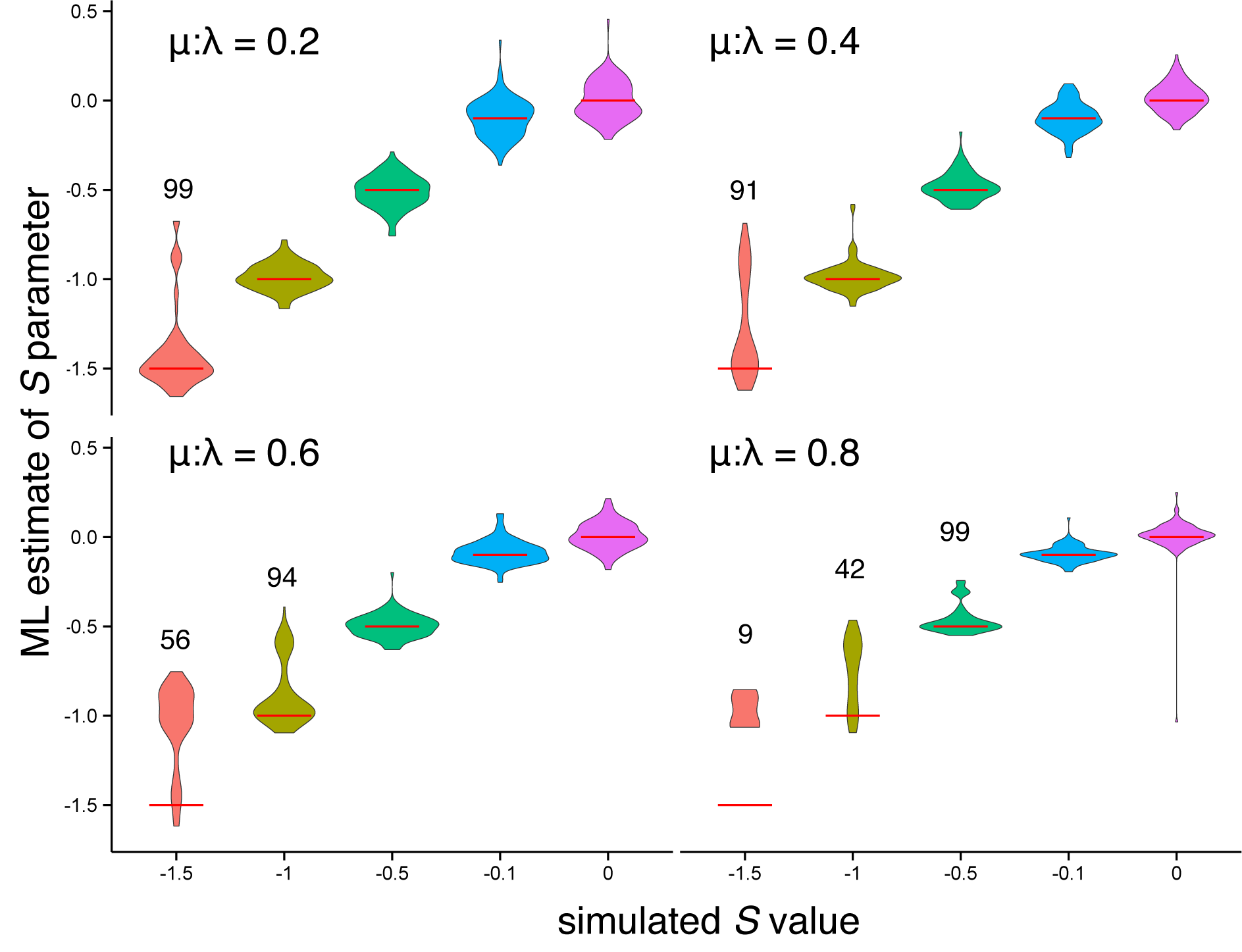
Simulation results showing the effect of varying the extinction fraction on estimation of the *S* parameter for the matching competition model. Red horizontal lines indicate the simulated *S* values, and numbers above sets of simulations indicate the sample size of included simulations under those scenarios (see main text for more details).

**Supplementary Figure 6.**
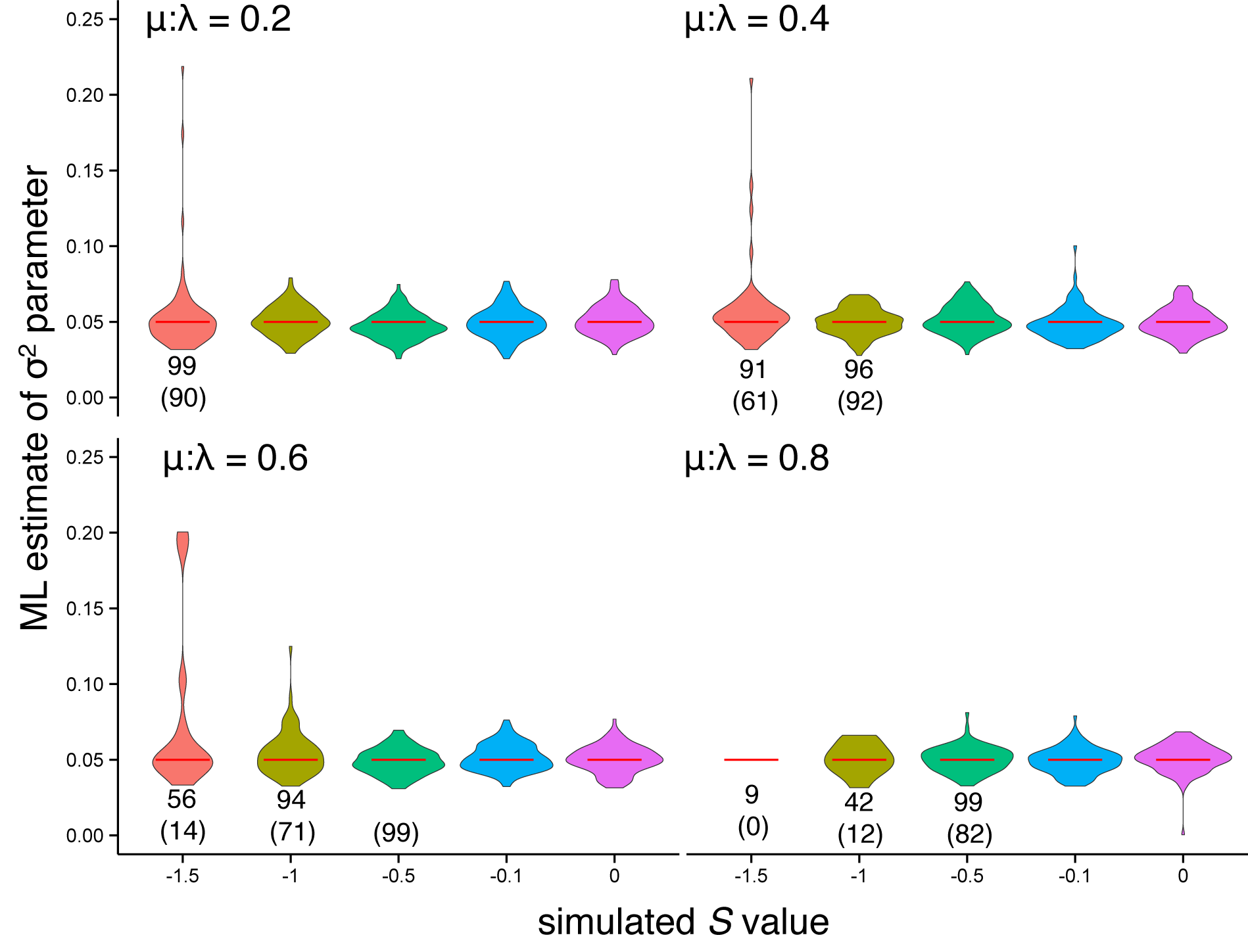
Simulation results showing the effect of varying the extinction fraction on estimation of the *σ*^2^ parameter for the matching competition model. Red horizontal lines indicate the simulated *σ*^2^ value (0.05), the numbers below sets of simulations indicate the sample size of included simulations under those scenarios (see main text for more details), and the number in parentheses indicate sample size after *σ*^2^ values > 0.25 were removed.

**Supplementary Figure 7.**
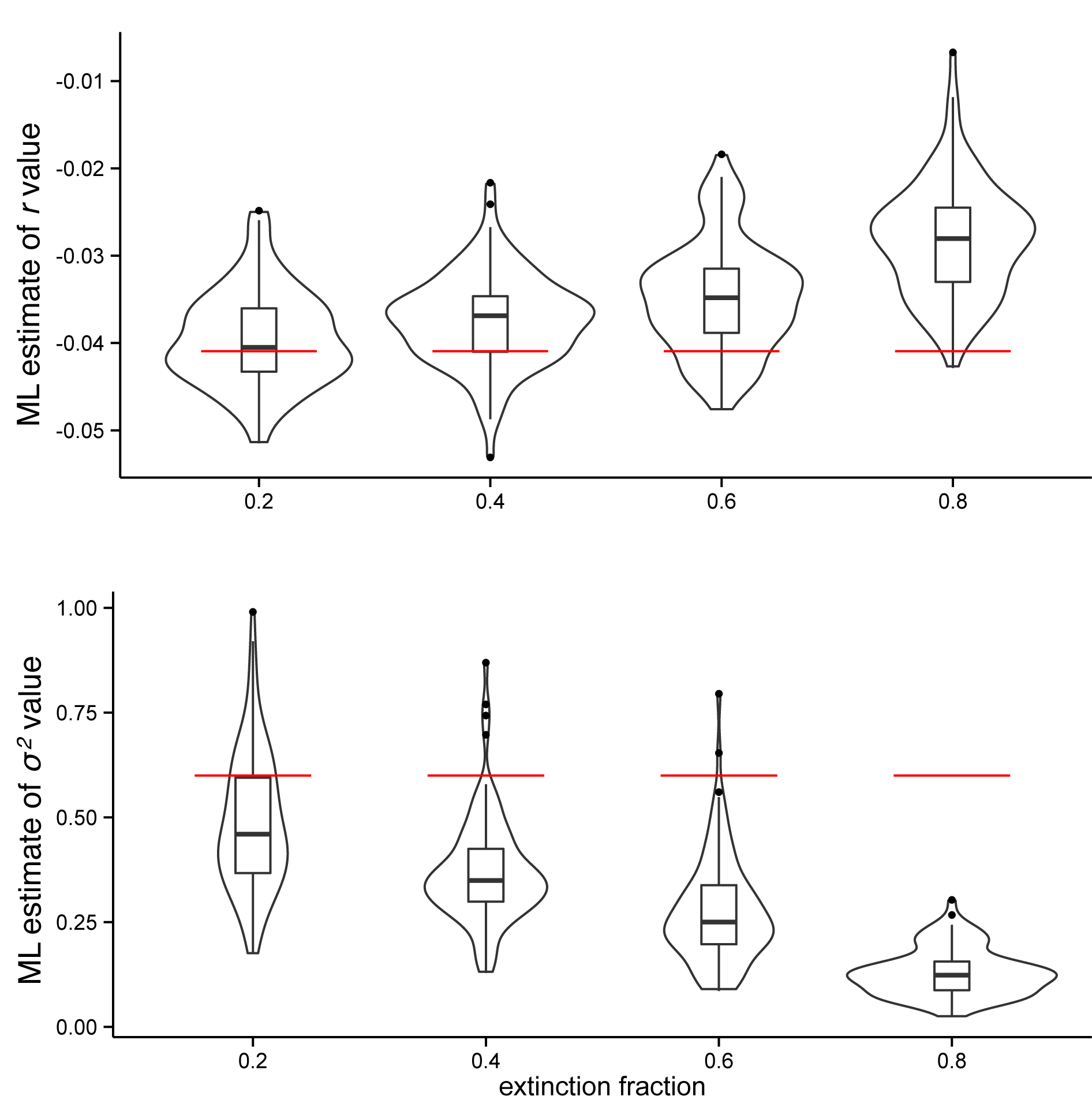
Simulation results showing the effect of varying the extinction fraction on slope (top) and *σ*^2^ (bottom) parameters for the exponential diversity-dependent model. Increasing extinction levels result in increasingly underestimated slope values and *σ*^2^ parameters. Red horizontal lines indicate the simulated parameter values.

**Supplementary Figure 8.**
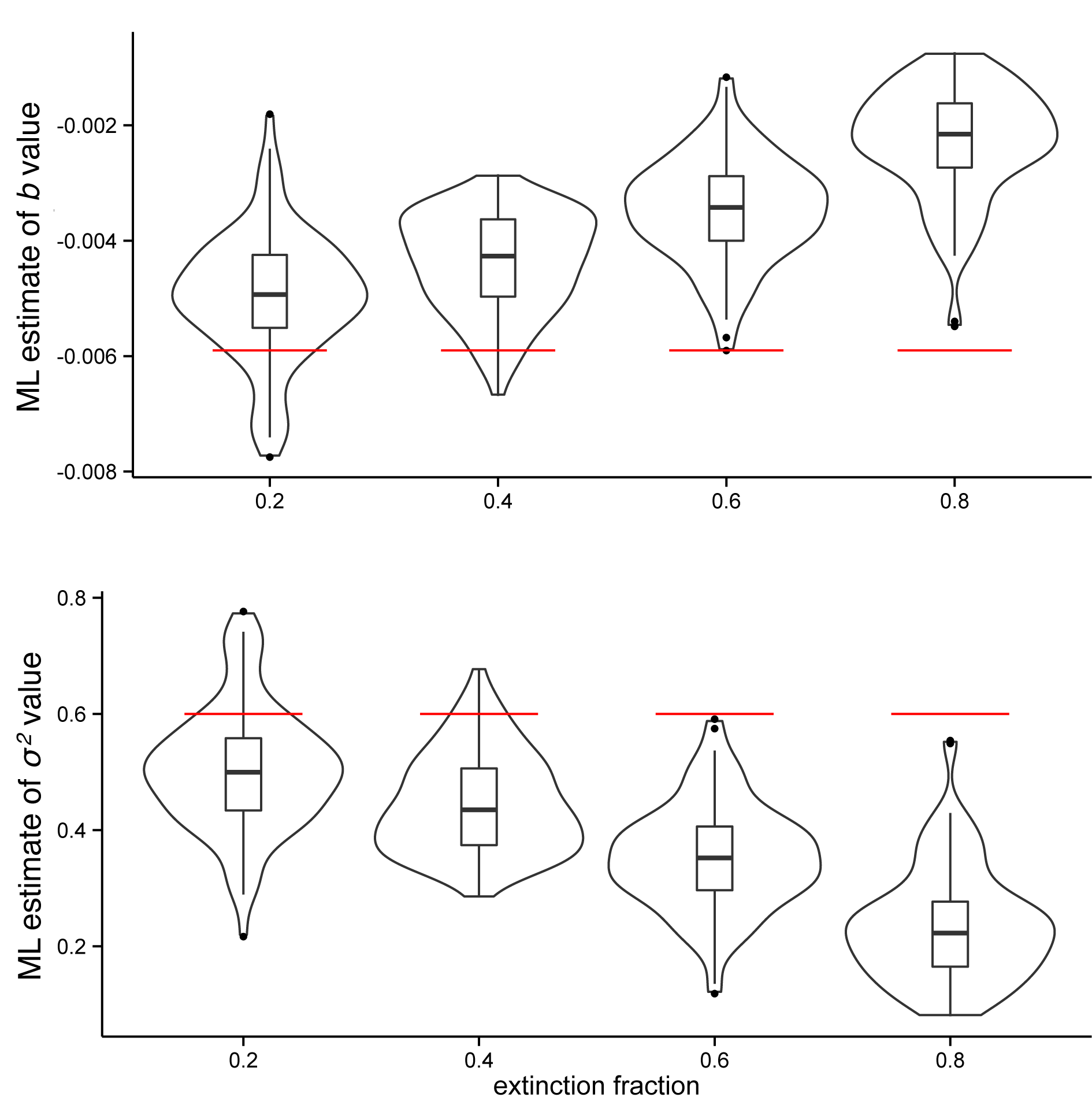
Simulation results showing the effect of varying the extinction fraction on slope (top) and sigma-squared (bottom) parameters for the linear diversity-dependent model. Increasing extinction levels result in increasingly underestimated slope values and *σ*^2^ parameters. Red horizontal lines indicate the simulated parameter values.

## 1 Supplementary Appendix 1

Considering that *n* lineages are interacting at time *t*, each trait *i* evolves following the equation:

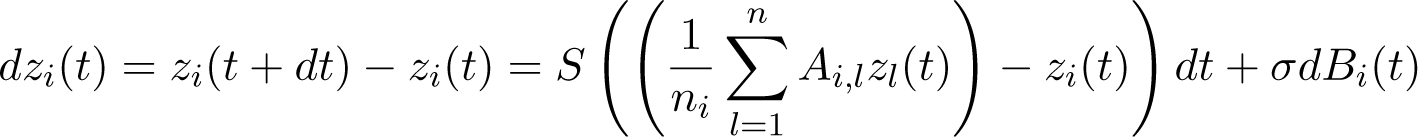

Where *A_i,l_* is equal to 1 if lineages *i* and *l* are sympatric, and to 0 otherwise, 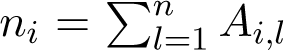 is the total number of lineages in sympatry with lineage *i*, and *B_i_* (*t*) represents standard Brownian motion.

Here, we present the derivation of Equations 3a and 3b from the main text. To make the derivation easier to follow, we drop the dependence on time *t*, replacing *z_i_* (*t*) with *z_i_* and *B_i_* (*t*) with *B_i_*.

First, applying the Itô formula to these stochastic processes gives us:

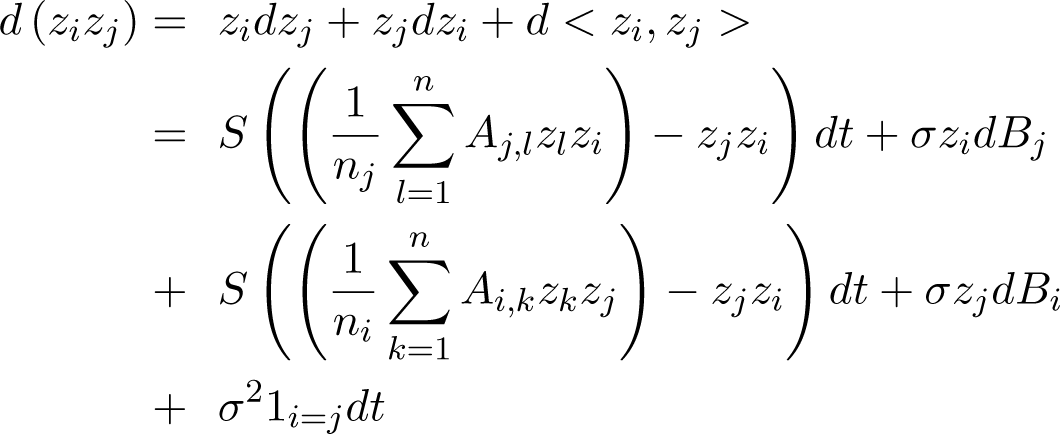

where 1_*i*=*j*_ equals one if *i* = *j* and zero otherwise.

Taking this expectation, it follows that:

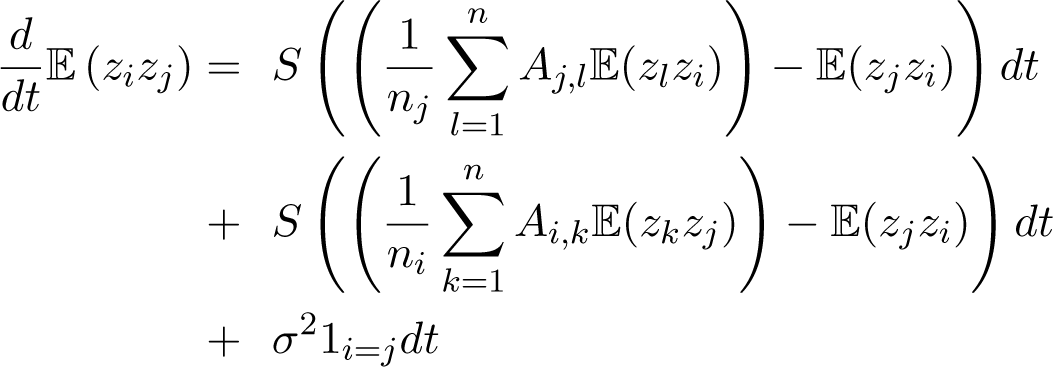

Moreover, we get:

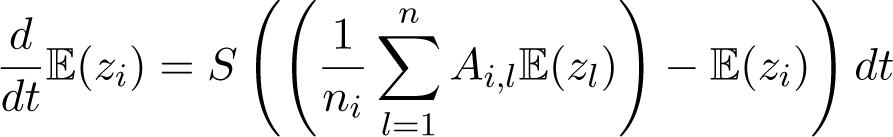

which leads to:

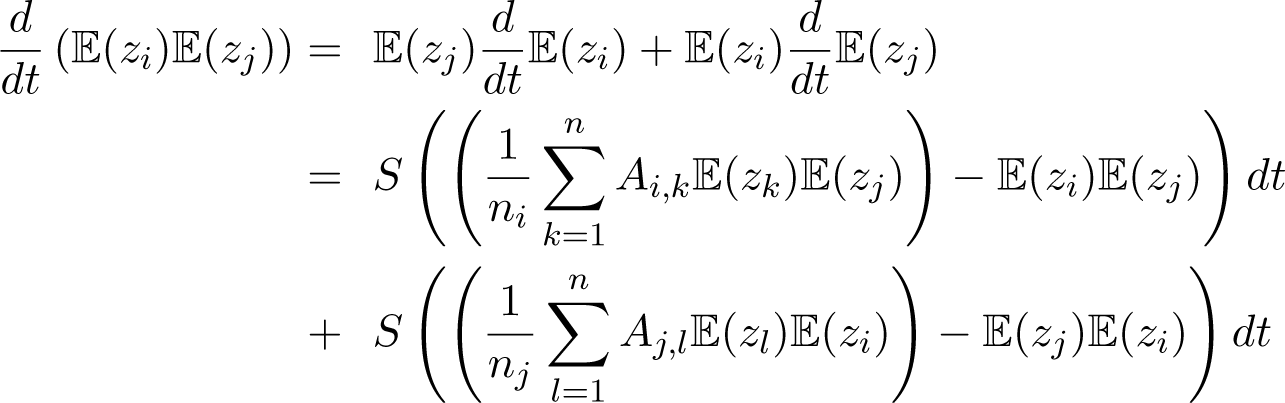

Taking together these different parts gives us the ODE satisfied by all covariances (denoted *v_i,j_* = Cov(*z_i_, z_j_*)):

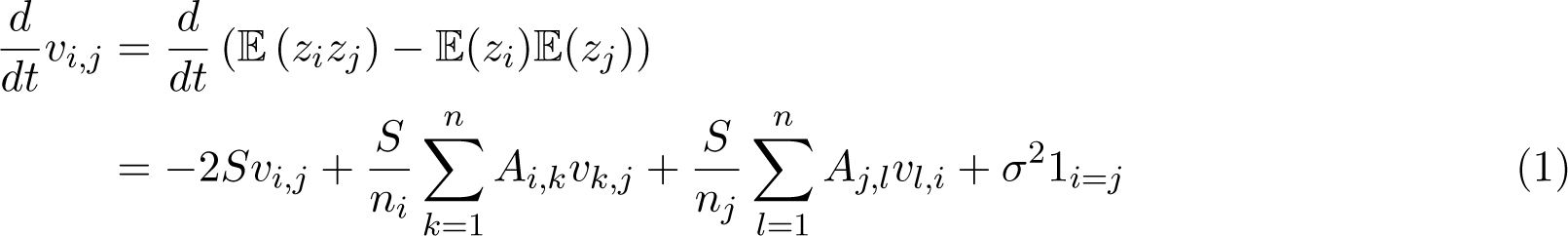

Using these derivations, the variance terms (*i* = *j*) are calculated using:

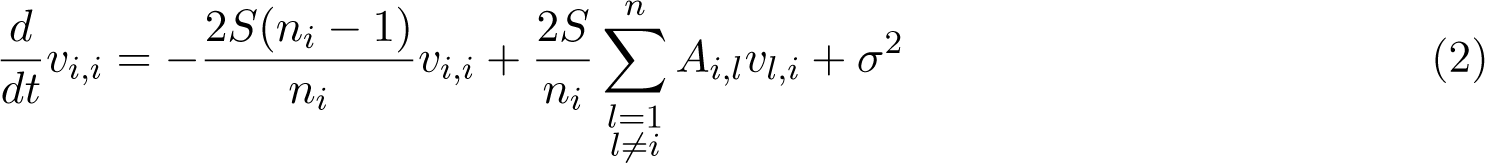

The covariance terms (*i* ≠ *j*) are calculated using:

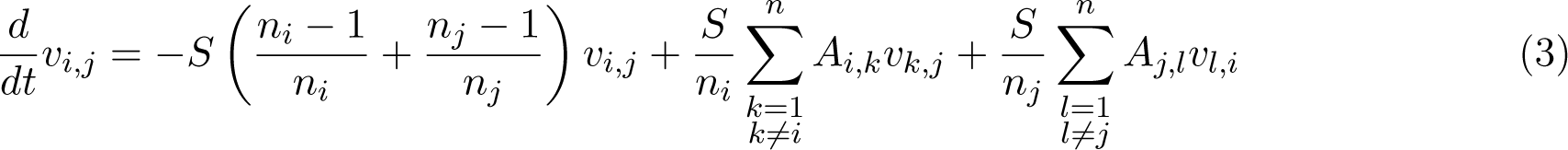

In the case where lineages *i* and *j* are in sympatry, this formula simplifies to:

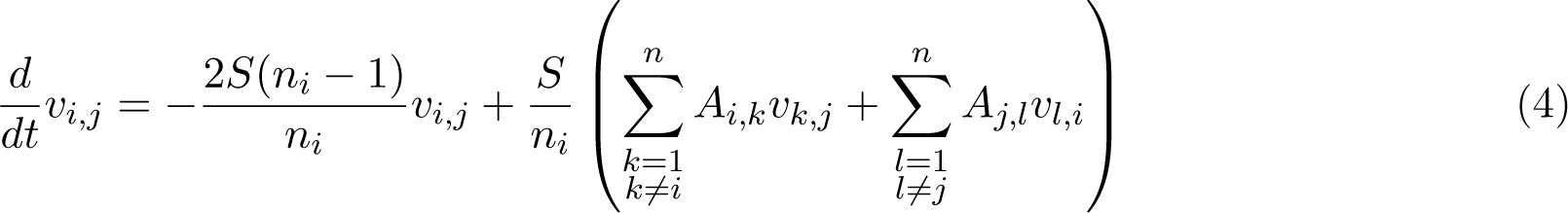

To solve the ODEs for the variance and covariance terms from the root to the tip, we begin by fixing the variance *v*_0_ for the process at the root to 0. At each speciation event, the starting value for both the variance of each of the new lineages and the covariance between the two new lineages is the variance of the immediate ancestor at the time of the speciation event, and the starting value for the covariance between the new lineages and any other persisting lineage is set to the value of the covariance between the persisting lineage and the ancestor of the new lineages at the time of speciation.

